# Genome-wide analysis of TFIIB’s role in termination of transcription

**DOI:** 10.1101/2024.02.22.581640

**Authors:** Michael J. O’Brien, Jared Schrader, Athar Ansari

**Affiliations:** Department of Biological Science, 5047 Gullen Mall, Wayne State University, Detroit, MI 48202

**Keywords:** Budding Yeast, Genome-wide Analyses, RNA polymerase II, TFIIB, Transcription

## Abstract

Apart from its well-established role in initiation of transcription, the general transcription factor TFIIB has been implicated in the termination step as well. The ubiquity of TFIIB involvement in termination as well as mechanistic details of its termination function, however, remains largely unexplored. To determine the prevalence of TFIIB’s role in termination, we performed GRO-seq analyses in *sua7-1* mutant (TFIIB^*sua7-1*^) and the isogenic wild type (TFIIB^*WT*^) strains of yeast. Almost a three-fold increase in readthrough of the poly(A)-termination signal was observed in TFIIB^*sua7-1*^ mutant compared to the TFIIB^*WT*^ cells. Of all genes analyzed in this study, nearly 74% genes exhibited a statistically significant increase in terminator readthrough in the mutant. To gain an understanding of the mechanistic basis of TFIIB involvement in termination, we performed mass spectrometry of TFIIB, affinity purified from chromatin and soluble cellular fractions, from TFIIB^*sua7-1*^ and TFIIB^*WT*^ cells. TFIIB purified from the chromatin fraction of TFIIB^*WT*^ cells exhibited significant enrichment of CF1A and Rat1 termination complexes. There was, however, a drastic decrease in TFIIB interaction with both CF1A and Rat1 termination complexes in TFIIB^*sua7-1*^ mutant. ChIP assay revealed that the recruitment of Pta1 subunit of CPF complex, Rna15 subunit of CF1 complex and Rat1 subunit of Rat1 complex registered nearly 90% decline in the mutant over wild type cells. The overall conclusion of these results is that TFIIB affects termination of transcription on a genome-wide scale, and TFIIB-termination factor interaction may play a crucial role in the process.

## Introduction

TFIIB is a general transcription factor (GTF) and an essential component of the preinitiation complex (PIC). During initiation of transcription, TFIIB performs a critical role in promoter recognition, RNAPII recruitment, and transcription start site selection (Woychik and Hampsey, 2002; Luse, 2014). It binds directly to sequences in the promoter region and makes multiple contacts with the components of the PIC (Lagrange et al., 1998; Deng and Roberts, 2005; Rhee and Pugh, 2012). Genome-wide analyses employing ChIP-Seq demonstrated TFIIB occupancy in the 5’ end of genes (Venters and Pugh, 2013; Pugh and Venters, 2016; Rossi et al., 2021) in accordance with its established role in initiation of transcription. It was, however, surprising to find TFIIB occupying the 3’ end of many genes in budding yeast (Singh and Hampsey, 2007; Mavrich et al., 2008; El Kaderi et al., 2009; Mayer et al., 2010; Medler et al., 2011; Murray et al., 2012; Al-Husini et al., 2013). At least in some genes, TFIIB ChIP signal at the 3’ end may be due to the presence of anti-sense promoter there (Mapendano et al., 2010). Another study using the ChIP-exo approach, however, failed to identify TFIIB at the 3’ end of yeast genes (Rhee and Pugh, 2012). The ChIP-exo approach involves digestion of immunoprecipitated chromatin with λ exonuclease, which digests the DNA except the region protected by the bound protein. The lack of 3’ TFIIB occupancy by ChIP-exo, therefore, suggests TFIIB does not directly bind to the DNA near the 3’ end.

The ChIP-exo results suggest that TFIIB crosslinks to the 3’ end of genes due to protein-protein interactions. This view is supported by the discovery of multiple interactions of TFIIB with the factors operating at the 3’ end of genes (reviewed in Al-Husini et al., 2020). One of the first and noteworthy among these interactions was with Ssu72, which is a subunit of the CPF 3’ end processing-termination complex. TFIIB exhibits both a genetic and physical interaction with Ssu72 (Pinto et al. 1994; Sun and Hampsey 1996; Wu et al., 1999; Allepuz-Fuster et al., 2019). Further investigation revealed interaction of TFIIB with yeast CF1 termination complex subunit Rna15 as well as its human homolog CstF64 (El Kaderi et al., 2009; Wang et al., 2010). The yeast study culminated in purification of a complex of TFIIB with Rna14, Rna15, Pcf11, Clp1 and Hrp1 subunits of CF1 complex (Medler et al., 2011). In addition to the CF1 complex, TFIIB was shown to interact with a number of subunits of CPF 3’ end processing-termination complex (Chereji et al., 2017). Recently, we demonstrated that the interaction of TFIIB with all three yeast termination complexes takes place almost exclusively in the context of transcriptionally active chromatin (O’Brien and Ansari, 2024). Taken together, these studies strongly suggested that TFIIB may have a hitherto unidentified role in the termination step of transcription.

Although most of the TFIIB-termination factor interactions were observed in yeast, the first report of the factor playing a role in termination came from a mammalian study (Wang et al., 2010). This study also found that TFIIB phosphorylation at serine-65 is necessary for its termination function. TFIIB serine-65 phosphorylation regulates its interaction with the Cstf-64 subunit of the CstF termination complex and facilitates recruitment of the complex at the 3’ end of a gene (Wang et al., 2010). In budding yeast, TFIIB’s involvement in termination was demonstrated using the *sua7-1* mutant, which has glutamic acid replaced by lysine at the 62^nd^ position (Allepuz-Fuster et al., 2019). In this mutant, the promoter recruitment of TFIIB remains unaffected, but its terminator occupancy is diminished (Singh and Hampsey, 2007). The interaction of TFIIB with the termination factors is also disrupted in the *sua7-1* mutant (Medler et al., 2011). The net result is that the recruitment of termination factors at the 3’ end of genes is compromised in the *sua7-1* mutant leading to a termination defect (Allepuz-Fuster et al., 2019). TFIIB also affects poly(A)-independent termination, which is an alternate pathway for termination of transcription unique to budding yeast (Roy and Chanfreau, 2018). A role of TFIIB in poly(A)-dependent termination was demonstrated in flies as well (Henriques et al., 2012).

TFIIB has been implicated in termination of transcription of just a few genes in budding yeast and humans, and only one gene in flies. The global role of TFIIB in termination, however, remains largely unexplored (Wang et al., 2010; Henriques et al., 2012; Allepuz-Fuster et al., 2019). The biochemical basis of TFIIB’s contribution to termination is also unclear. Understanding the prevalence of TFIIB involvement in termination and the mechanistic basis of this function is crucial for understanding its comprehensive role in the transcription cycle. We therefore performed GRO-seq analysis and mass spectrometry of affinity purified TFIIB from wild type and the *sua7-1* mutant. Our results demonstrate that TFIIB-termination factor interaction affects termination of transcription on a genome-wide scale in budding yeast.

## Results

### TFIIB affects termination of transcription on a genome-wide scale

To determine the role of TFIIB in termination, we used the *sua7-1* mutant of TFIIB (TFIIB^*sua7-1*^). This mutant has glutamic acid replaced by lysine at position 62 (E62K) (Pinto et al., 1994). The mutant is cold-sensitive and exhibits altered transcription start site selection (Sun and Hampsey, 1996). The binding affinity of TFIIB^*sua7-1*^ for the promoter, as well as its interactions with TBP and RNAPII, are comparable to that of wild type TFIIB (TFIIB^*WT*^) (Cho and Buratowski, 1999). The recruitment of general transcription factors and RNAPII onto the promoter during assembly of the preinitiation complex is also indistinguishable from wild type (Cho and Buratowski, 1999). TFIIB has been observed to crosslink to the 3’ end of genes in TFIIB^*WT*^ cells (Singh and Hampsey, 2007). In TFIIB^*sua7-1*^, however, the 3’ end crosslinking is lost and consequently, promoter-terminator crosstalk or gene looping is diminished as well (Singh and Hampsey, 2007). Therefore, the TFIIB^*sua7-1*^ mutant is a suitable target for investigating a termination defect at the 3’ end of genes.

To determine the role of TFIIB in termination, we performed ‘<u>T</u>ranscription <u>R</u>un-<u>O</u>n’ (TRO) assays in TFIIB^*sua7-1*^ and TFIIB^*WT*^ cells for *BLM10, HEM3, SEN1, CBK1, KAP123* and *SUR1* genes. These genes were selected because their next immediate downstream gene is at least 500 bp away and therefore any termination defect can be determined with confidence. The TRO assay detects the presence of transcriptionally active polymerases on a gene. When termination is defective, TRO signal is detected downstream of the poly(A) termination site (Figure 1A). Strand-specific TRO analysis revealed that in all six genes, there was a strong polymerase signal in the coding region before the poly(A) site in wild type cells (Figure 1B and 1C, blue bars). There was, however, a strongly reduced or undetectable polymerase signal in the region downstream of the poly(A) site (Figure 1B and 1C, blue bars). In TFIIB^*sua7-1*^ cells, though there was no difference in the TRO signal upstream of the poly(A) site compared to the wild type, there was a dramatic increase in TRO signal in the downstream regions of all six genes (Figure 1B and 1C, red bars). This revealed failure to read the termination signals efficiently in the TFIIB^*sua7-1*^ mutant, and continued transcription of downstream regions. The termination defect in TFIIB^*sua7-1*^ cells is not due to an indirect effect of mutation on termination factor levels as Western blot signal of Rna15 and Pta1 is similar in the wild type and TFIIB^*sua7-1*^ cells (Supplementary Figure 1).

**Figure 1:**
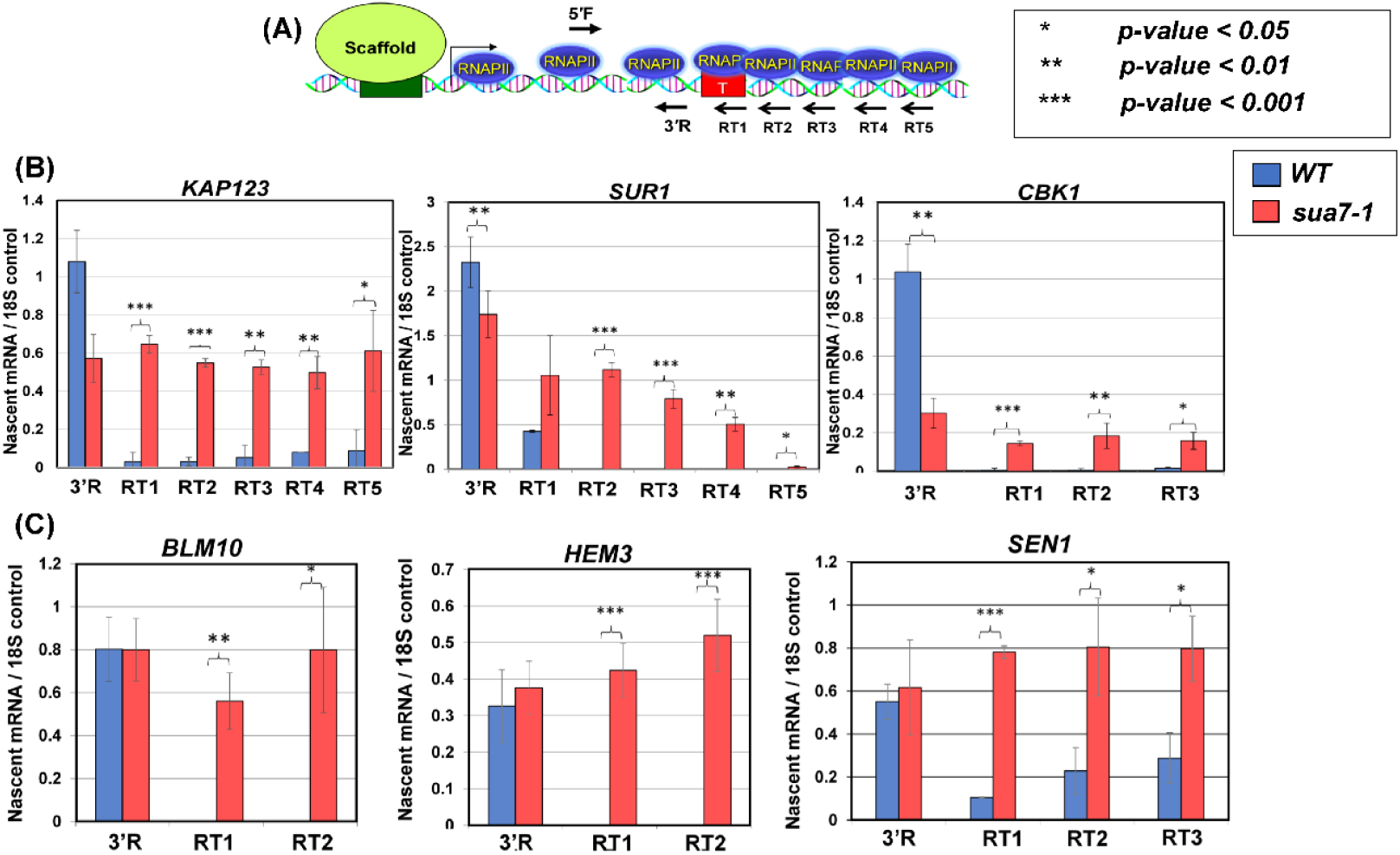
TRO shows a transcription termination defect for six genes in the sua7-1 mutant. (A) Schematic depiction of a gene showing the actively transcribing RNA polymerase II and positions of primers used for cDNA synthesis following TRO procedure. Primers are depicted with arrows; primers within the gene body are primers 5’F and 3’R whereas primers RT1-5 are downstream of the terminator region. **(B, C)** Quantification of RNA levels detected following TRO analysis in in wild type and sua7-1 mutant cells. Results shown here are from three biological replicates. RT-PCR signal is represented as nascent mRNA signal compared to 18S control. RT reactions performed with individual primers as shown in diagram. PCR reaction performed with primer pair 5’F/3’R. p-values are a result of a standard t-test. Error bars represent standard deviation.

These results prompted us to examine TFIIB’s involvement in termination on a genome-wide scale. To accomplish this we used the ‘<u>G</u>lobal <u>R</u>un-<u>O</u>n-Seq’ (GRO-seq) approach, which is the genome-wide version of the strand-specific TRO approach (Core et al., 2008). Briefly, GRO-seq involves incorporation of BrUTP in newly synthesized RNA, affinity purification of nascent labelled-RNA, reverse transcription, and high throughput sequencing of cDNA library as shown in Figure 2A. GRO-seq provides a high resolution snapshot of the position and density of actively transcribing polymerase in a strand specific manner (Figure 2B). GRO-seq was performed with TFIIB^*sua7-1*^ and TFIIB^*WT*^ strains with three biological replicates. GRO-seq samples were first stripped of adapter sequences using cutadapt. The 3’ end reads were then aligned to the yeast s288c genome downloaded from SGD (version R64-3-1). Throughout this analysis, we used the 3’ annotated end of mRNAs from SGD for cells grown in YPD media corresponding to the poly(A) cleavage site. We restricted our analysis to 2501 protein coding genes whose 3’ ends were at least 500 bp away from the neighboring gene on the same strand. This was done because of the compact nature of the yeast genome, which often results in the terminator region of a gene overlapping with the promoter or terminator elements of the neighboring gene. To restrict our analysis to transcriptionally active genes in YPD growth conditions, we selected mRNAs that were at least 500 nucleotides in length or longer, and whose expression value was more than or equal to 1 read/nucleotide within the CDS in at least three biological replicates. We therefore performed our final analysis with 337 genes that satisfied all benchmarks described above. To avoid the results being skewed in favor of long and highly expressed genes, we normalized by dividing the number of reads with the length of the gene.

**Figure 2:**
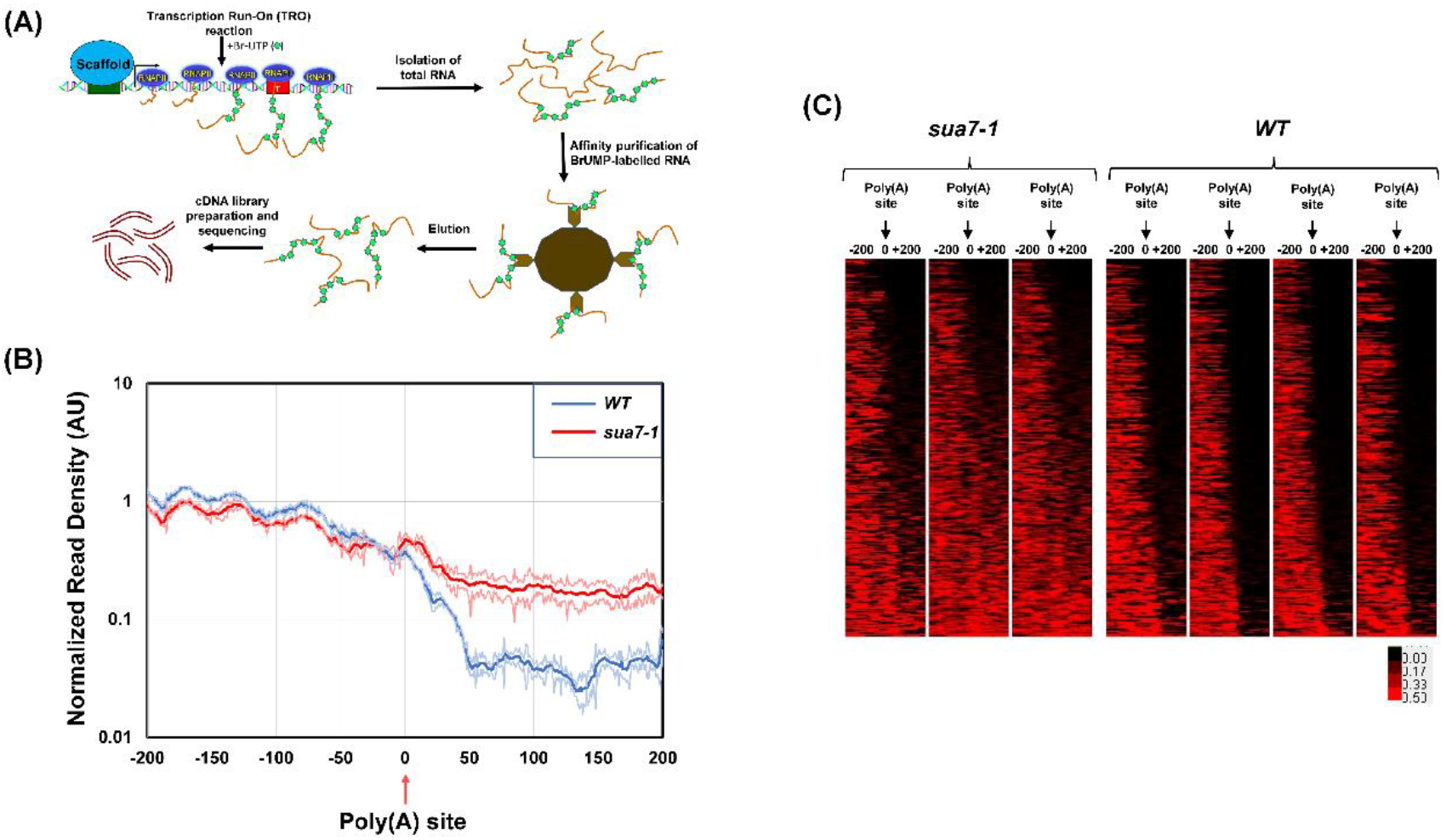
GRO-seq demonstrates a genome-wide defect in transcription termination in the sua7-1 mutant. **(A)** Schematic showing the major steps of the GRO-seq approach, culminating in a sample prepared for sequencing. (**B)** Metagene plot of normalized read densities. WT and sua7-1 normalized read densities are plotted from -200 nt upstream of poly(A) site to +200 nt downstream of poly(A) site. Thick/center lines are the average normalized read densities. Upper and lower (light colored) lines represent standard error. Data is from three biological replicates. **(C)** Heat map shows GRO-seq signal of the same set of 337 genes in all three replicates arranged from top to bottom in order of increasing terminator readthrough phenotype in TFIIB^sua7-1^ cells. For each gene, the change in GRO-seq signal in TFIIB^sua7-1^ and TFIIB^WT^ cells is indicated according to the red/black scale.

To compare the termination of transcription in TFIIB^*sua7-1*^ and TFIIB^*WT*^ cells, GRO-seq reads were aligned in a 200 bp window upstream and 200 bp downstream of the poly(A) site. A metagene plot shows both TFIIB^*WT*^ and TFIIB^*sua7-1*^ exhibiting a similar GRO-seq profile in the region upstream of the poly(A) site (Figure 2B). There was, however, a sharp drop in number of reads beyond the poly(A) site in TFIIB^*WT*^ cells (Figure 2B, blue line). In TFIIB^*sua7-1*^ also there was a decrease in the number of reads after the poly(A) site, but the number of reads was higher than in TFIIB^*WT*^ (Figure 2B, red line). Heat maps of individual replicates demonstrate that there is hardly any detectable GRO-seq signal beyond the poly(A) site in TFIIB^*WT*^ cells in all replicates (Figure 2C). In the TFIIB^*sua7-1*^ mutant, however, nearly 70% of genes exhibited a varying degree of GRO-seq signal in the region downstream of the poly(A) site in all three replicates (Figure 2C). Genes in the bottom 40% section of the heat map show strong GRO-seq signal while upper 25% of genes show no GRO-seq signal after the poly(A) site in the mutant (Figure 2C). These results indicate that, in nearly two-thirds of the analyzed genes, the polymerase was unable to read the poly(A) termination signal efficiently in TFIIB^*sua7-1*^ cells leading to an enhanced readthrough in the region downstream of the poly(A) site (Figures 2C).

To further characterize the role of TFIIB in termination on a genome-wide scale, we calculated the readthrough index (RTI) in the mutant and wild type cells as described in (Baejen et al., 2017). The RTI was calculated by measuring the ratio between the number of read counts 500 bp downstream and 500 bp upstream of the poly(A) site, excluding a region 50 bp directly up- and downstream of the poly(A) site as shown in Figure 3A. A termination defect is expected to result in more GRO-seq reads downstream of the poly(A) site compared to the upstream region and therefore a higher RTI value.

**Figure 3:**
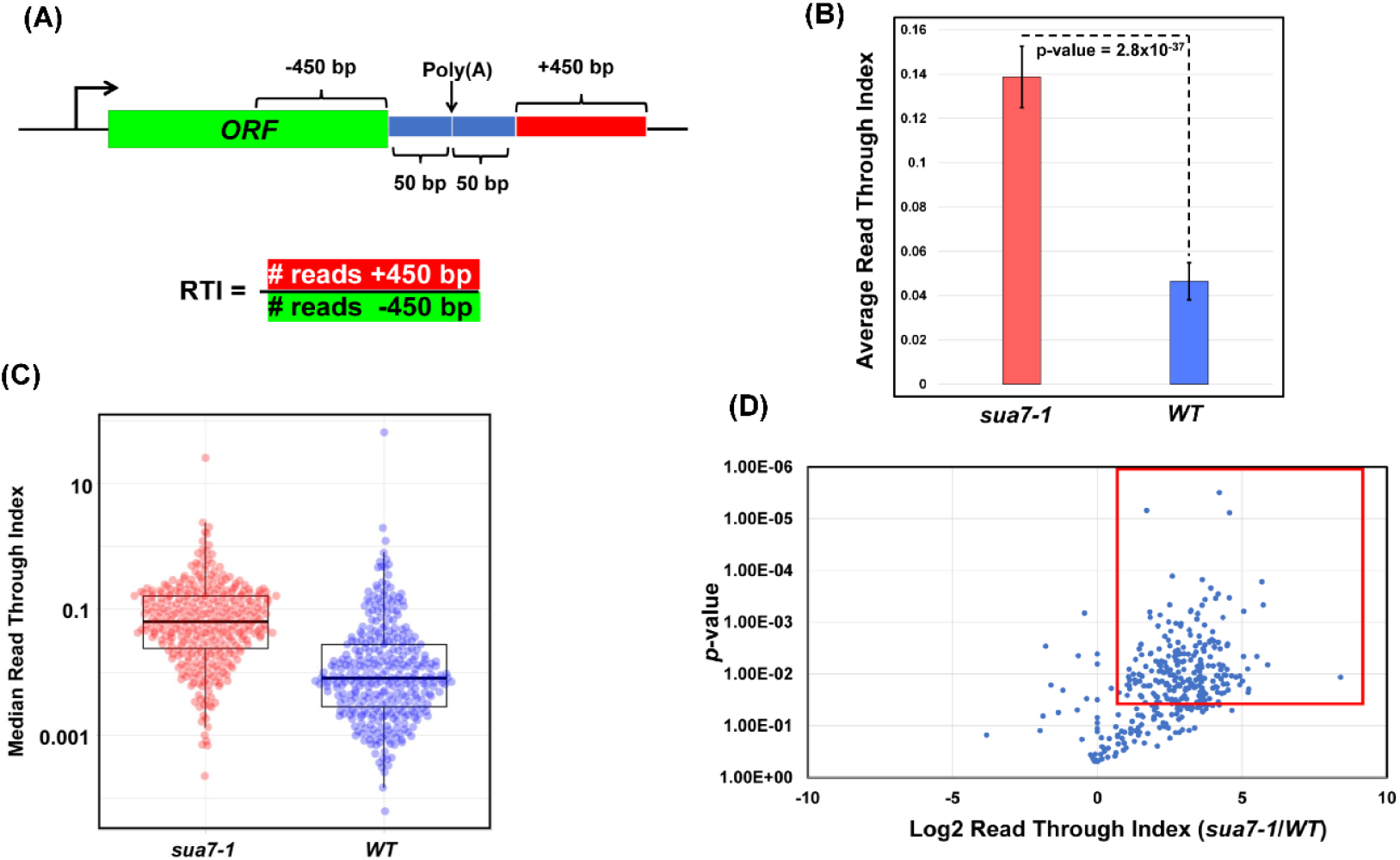
A majority of genes have an increased Read Through Index in sua7-1. **(A)** Schematic representing calculation of Read Through Index (RTI) for individual genes. RTI is calculated from number of reads +450 bp divided by number of reads -450 bp. **(B)** The average Read Through Index of all analyzed genes in sua7-1 and WT cells. Data represented is from three biological replicates and p-value obtained from standard 2-sided, paired t-test. **(C)** Median Read Through Index for all analyzed genes in sua7-1 and WT cells represented as a box and whisker plot. The middle line represents the median value of the corresponding RTI. The top and bottom lines represent the upper and lower quartiles. Data is represented from three biological replicates and p-value obtained from standard 2-sided t-test with unequal variance. n=337 is the total number of analyzed genes. **(D)** The log2 RTI ratio of sua7-1/WT is plotted on the x-axis with p-value plotted on the y-axis. All genes analyzed are plotted as blue dots. The outlined red window contains all genes that display at least a 2-fold increase in RTI ratio of sua7-1/WT while simultaneously having a p-value less than 0.05. Data is represented from three biological replicates and p-value obtained from standard 1-sided t-test with unequal variance.

The RTI value in the mutant (RTI=0.139) registered a statistically significant 3-fold increase over the wild type cells (RTI=0.046) (Figure 3B). These results suggest that TFIIB affects termination of transcription on a genome-wide scale in budding yeast. The RTI values in Figure 3B represent mean values, which are a good indicator of the trend if data follow a symmetric distribution. The mean values, however, can be skewed by outliers. In contrast, median values are not affected by outliers and more accurately represent the general trend. Therefore, to further probe if termination in the mutant is affected on a genome-wide scale, median RTI values of genes in the mutant and the wild type strains for all replicates were charted in a box and whisker plot. Median RTI values calculated from this plot are 0.063 and 0.008 for TFIIB^*sua7-1*^ and TFIIB^*WT*^ respectively (Figure 3C), making the median RTI >7-fold higher than wild type. These results corroborate the conclusion drawn from mean RTI values shown in Figure 3B that TFIIB affects termination of transcription on a genome-wide scale.

We next determined what fraction of genes used in this analysis display defective termination in TFIIB^*sua7-1*^. The log2 values of RTI ratios of TFIIB^*sua7-1*^ / TFIIB^*WT*^ for all genes were plotted as a function of *p*-values (Figure 3D). The boxed area in the plot represents all genes that exhibit log2 RTI ratio of more than one, indicative of a 2-fold increase in the RTI of TFIIB^*sua7-1*^ compared to TFIIB^*WT*^, and a *p*-value of less than 0.05. Based on these criteria, nearly 73.8% genes analyzed in this study show a statistically significant dependence on TFIIB for efficient termination of their transcription. This is in contrast to Pcf11 and Ysh1, which affect termination of more than 90% of genes in yeast (Baejen et al., 2017). Thus, TFIIB is not an essential termination factor like CPF and CF1 3’ end processing-termination complexes but has a significant effect on termination of nearly three fourths of genes used in our analysis.

Next, we searched for common features among genes that conferred dependence on TFIIB for efficient termination of transcription. Genes whose termination is affected by TFIIB do not display common predicted structures or enriched sequence motifs. TFIIB-linked termination is also not related to the length of genes (Supplementary Figure 2A). Furthermore, the TFIIB-associated termination phenotype could not be linked to the presence or absence of TATA-box in the promoter of the gene (Supplementary Figure 2B). Both TATA and TATA-less promoter containing genes exhibited similar RTI value distribution in TFIIB^*sua7-1*^ strain (Supplementary Figure 2B). Ontological analysis revealed that TFIIB termination dependence was more pronounced in genes associated with translation and ribosomal function which is not surprising as the fraction analyzed here was already enriched in genes associated with ribosomal function (Supplementary Figure 2C). Thus, TFIIB role in termination cannot be attributed to a particular functional category of genes.

### TFIIB’s interaction with termination factors is diminished in the termination defective mutant

Having established the role of TFIIB in termination of transcription of nearly three-fourths of analyzed genes in budding yeast, we next investigated the mechanism underlying the role of the factor in termination. We hypothesized that TFIIB may affect termination directly by interacting with the termination factors and regulating their activity at the 3’ end of genes or indirectly by affecting CTD-serine-2 phosphorylation which in turn facilitates the recruitment of termination factors. Our recently published results favored the first possibility (O’Brien and Ansari, 2024). We demonstrated interaction of TFIIB with all three termination complexes of yeast; CPF, CF1 and Rat1, in the TFIIB^*WT*^ strain. The interaction with CF1 and Rat1 complexes was observed exclusively in the chromatin context, thereby suggesting that the interaction occurred only during transcription. We reasoned that if the TFIIB-termination factor interaction is linked to a TFIIB-mediated effect on termination, then interaction of TFIIB with the termination factors will be reduced in the termination defective TFIIB^*sua7-1*^ strain. We therefore affinity purified TFIIB from TFIIB^*sua7-1*^ cells. Purification was performed from both chromatin and soluble fractions as described in O’Brien and Ansari (2024). Affinity purified fractions were subjected to mass spectrometry. A quantitative proteomic approach was followed to analyze mass spectrometry data (Paoletti et al., 2006; Zybailov et al., 2006). The relative abundance of a termination factor in a purified TFIIB sample was quantified by dividing the SAF value of the factor with that of TFIIB to get the TFIIB-normalized spectral abundance factor (BNSAF). The data presented here is the result of four independent replicates. Comparison of the BNSAF values revealed that the interaction of TFIIB with Rna14, Rna15, Pcf11 and Clp1 subunits of CF1 complex as well as Rat1 and Rai1 subunits of Rat1 complex exhibited a statistically significant decline of about 70-90% in TFIIB^*sua7-1*^ compared to TFIIB^*WT*^ cells (Figure 4B). Interaction with subunits of CPF complex also registered a drop in the mutant, but it was not statistically significant due to higher sample-to-sample variability (Figure 4B). We next checked if interaction of TFIIB with GTFs is similarly negatively impacted in the mutant. Contrary to the expectation, TFIIB interaction with subunits of TFIID, TFIIA and TFIIF exhibited an increase in TFIIB^*sua7-1*^ compared to TFIIB^*WT*^ cells, but the difference in BNSAF value between TFIIB^*sua7-1*^ and TFIIB^*WT*^ cells was statistically not significant (Supplementary Figure 3). The termination defective TFIIB^*sua7-1*^ mutant therefore exhibits a selective diminution in interaction with termination factors. These results suggest that TFIIB either helps in the recruitment of termination factors or stabilizes their recruitment at the 3’ end of genes.

**Figure 4:**
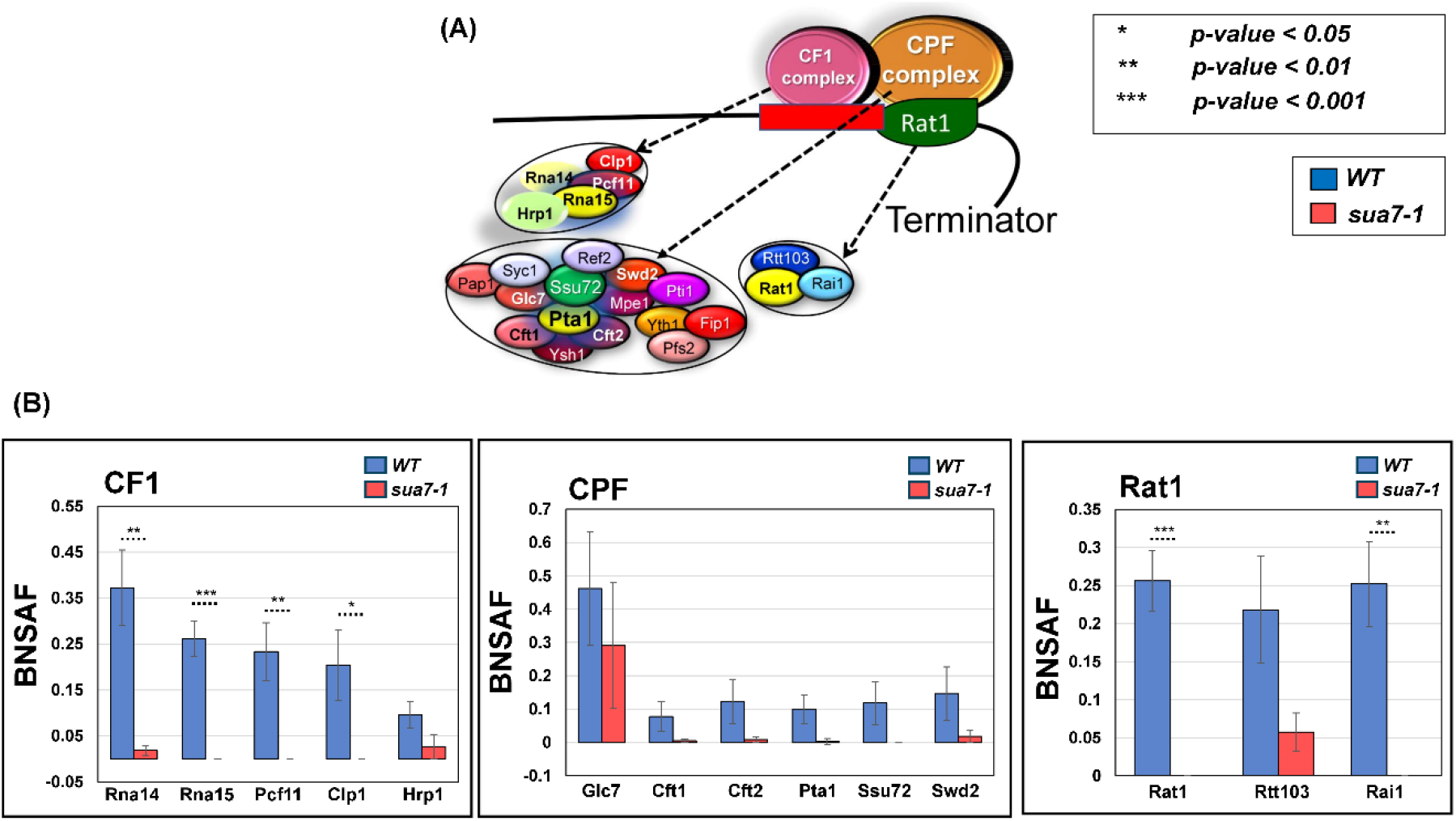
TFIIB-termination factor interactions are compromised in sua7-1 mutant. **(A)** Diagram of the three major termination complexes in S. cerevisiae and their corresponding subunits. **(B)** BNSAF values of termination factors, grouped by corresponding complexes in sua7-1 and WT cells. Data is from 4 biological replicates. p-values obtained from a standard, paired t-test. Error bars represent one unit of standard error based on four biological replicates.

### Recruitment of termination factors at the 3’ end of genes is compromised in the termination defective mutant

To further probe the role of the TFIIB-termination factor interaction in termination, we checked the recruitment of all three termination complexes at the 3’ end of *BLM10, HEM3, KAP123* and *SUR1*. These four genes were selected because their next neighboring gene on the same strand is more than 500 bp away and they exhibit a termination defect in TFIIB^*sua7-1*^ cells (Figure 1). The recruitment of the CPF complex was monitored in terms of its Pta1 subunit, of CF1 complex in terms of its Rna15 subunit and of Rat1 complex using its Rat1 subunit. All three subunits were tagged at their C-terminus by an HA-epitope and their recruitment at the 3’ end of *BLM10, HEM3, KAP123* and *SUR1* was examined in TFIIB^*sua7-1*^ and TFIIB^*WT*^ cells employing the ChIP approach. ChIP signal for each factor was normalized with the input signal and then with the RNAPII signal at the site. As expected, all three factors, Pta1, Rna15 and Rat1 were localized at the 3’ end of genes in TFIIB^*WT*^ cells (Figure 5B, 5C and 5D, blue bars). In TFIIB^*sua7-1*^ cells, however, the recruitment of all three factors registered a drop of nearly 90% (Figure 5B, 5C and 5D, red bars). The decrease in 3’ end crosslinking was statistically significant as the observed *p*-value in every case was less than 0.05 (Figure 5B, 5C and 5D). A possible interpretation of these results is that TFIIB is either required for the recruitment of the termination factors or for stabilization of their association with the 3’ end of genes or both. The possibility of TFIIB enhancing the exoribonuclease activity of Rat1 thereby expediting the dissociation of the elongating polymerase from the template, however, still cannot be ruled out.

**Figure 5:**
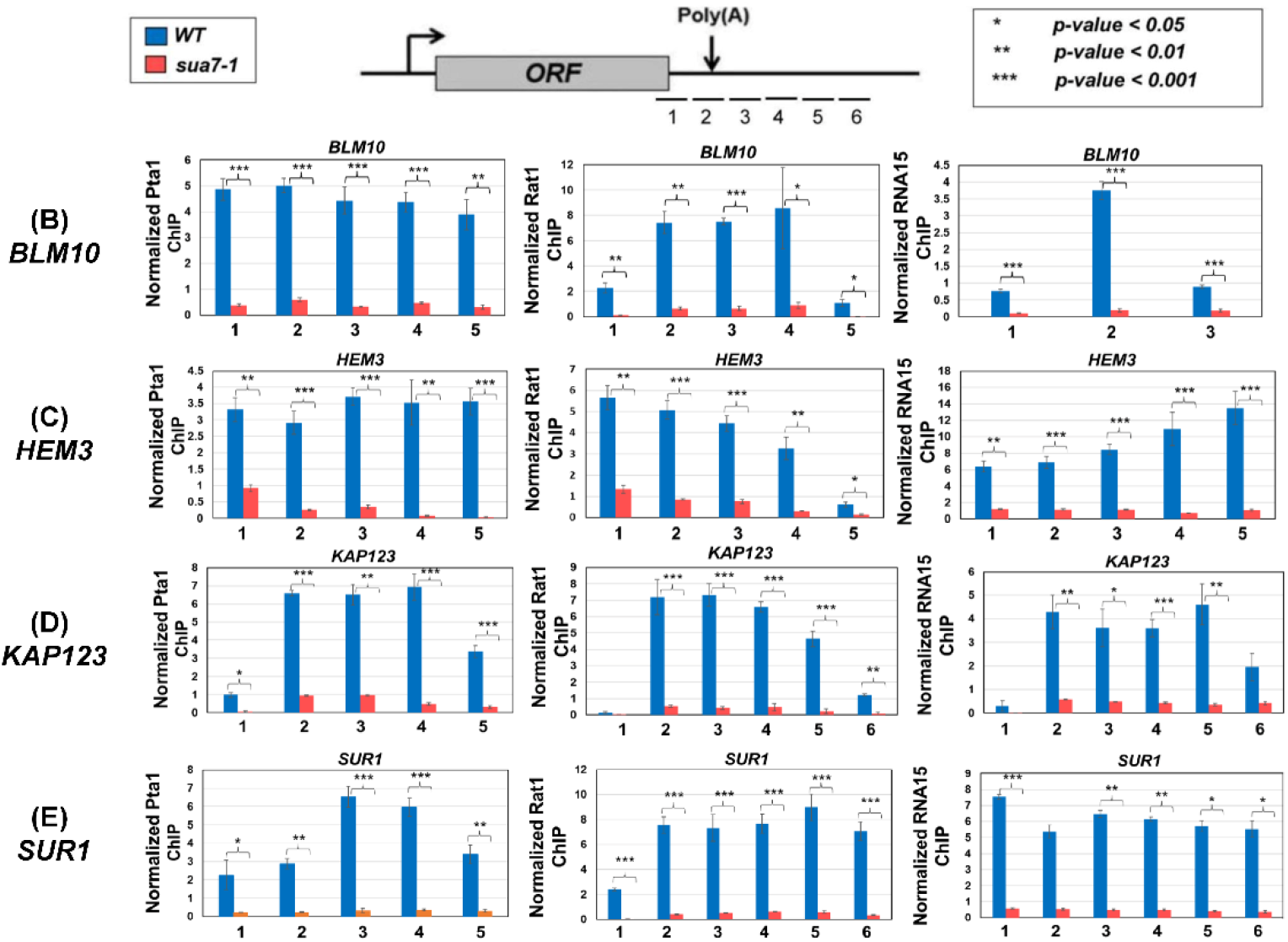
Termination factor occupancy is reduced at the 3’ end of genes in sua7-1 mutant. **(A)** Schematic depicting the primer locations for examining occupancy of termination factors on the terminator region and regions directly flanking the terminator in reference to a gene/ORF. **(B, C, D, E)** ChIP signal for each factor was normalized with the input signal and then with the RNAPII signal in the region. ChIP signal for Pta1, Rat1, and Rna15 at and around the terminator region is significantly decreased in sua7-1 compared to wild type cells for the genes BLM10, HEM3, KAP123, and SUR1. p-values were calculated by standard two-tailed t-test. Error bars represent one unit of standard error based on three biological replicates.

## Discussion

The role of the initiation step in transcription of a gene is well established. It is generally, however, not appreciated that the termination step is just as vital for efficient transcription of a gene. The generally accepted view is that transcription termination in budding yeast requires three multiprotein complexes; CF1, CPF and Rat1 complexes (Mischo and Proudfoot, 2013; Baejen et al., 2017). We show here that the general transcription factor TFIIB is also involved in termination of transcription on a genome-wide scale in budding yeast. Our conclusion is based on four crucial observations. First, TFIIB crosslinks to the 3’ end of a number of transcriptionally active genes in wild type cells but not in TFIIB^*sua7-1*^ mutant (Singh and Hampsey, 2007). Second, RNAPII reads through the termination signal in TFIIB^*sua7-1*^ mutant. The mutant exhibits a statistically significant increase of two-fold or more in RTI over the TFIIB^*WT*^ cells for at least 74% of genes analyzed in our study (Figure 2B). Third, TFIIB interacts with subunits of CF1 and Rat1 3’ end processing-termination complexes in wild type cells. In the termination defective TFIIB^*sua7-1*^ mutant, there was a statistically significant reduction in TFIIB interaction with termination factors (Figure 4), suggesting that the TFIIB-termination factor interaction may be crucial for efficient termination. Fourth, recruitment of Pta1, Rna15 and Rat1 termination factors at the 3’ end of genes registered a nearly 90% decline in TFIIB^*sua7-1*^ mutant over TFIIB^*WT*^ cells (Figure 5). The overall conclusion of these results is that TFIIB is involved in termination of transcription of nearly three-fourths of genes analyzed in this study in budding yeast.

TFIIB, however, is not an essential termination factor like Pcf11 and Ysh1 subunits of CF1A and CPF complexes respectively. In TFIIB^*sua7-1*^ mutant, there was no complete loss of termination as was observed upon nuclear depletion of Pcf11 and Ysh1 (Baejen et al., 2017). Despite the higher GRO-seq signal downstream of the poly(A)-site, the smooth descending polymerase signal was preserved in the mutant. A logical interpretation of these results is that termination still occurs in the mutant, but a significant number of polymerase molecules are unable to recognize the poly(A) signal and read through into the downstream region. These results are in agreement with the termination factors ChIP data that found 3’ end recruitment of all three termination complexes decreasing by about 90% in TFIIB^*sua7-1*^ mutant. A recent genome-wide analysis carried out in mammalian cells also reported polymerases being unable to read the termination signal in the absence of functional TFIIB in the cell (Santana et al., 2022). This study performed PRO-seq upon nuclear depletion of TFIIB and found a majority of polymerases reading through the termination signal. The extent of readthrough in the absence of TFIIB in this case was comparable to that observed upon nuclear depletion of Pcf11 and Ysh1 in yeast (Baejen et al., 2017). It is quite possible that the point mutation in TFIIB^*sua7-1*^ affects some aspects of termination, but the mutant TFIIB still retains interactions that allow it to facilitate termination, though with reduced efficiency. It will be interesting to look at the termination phenotype in yeast cells that are depleted of TFIIB in the nucleus by the anchor away approach. These results also demonstrate that the termination function of TFIIB has been conserved during evolution.

It is possible that the termination defect observed in TFIIB^*sua7-1*^ mutant is an indirect effect of the mutation on termination through defective initiation or elongation, and not due to the direct involvement of TFIIB in termination. The evidence, however, support TFIIB’s direct involvement. First, the mutant form of TFIIB is indistinguishable from the wild type counterpart in terms of its ability to initiate transcription *in vitro* (Cho and Buratowski, 1999). Second, the GRO-seq profile presented here shows almost identical signals in the open reading frame in TFIIB^*WT*^ and TFIIB^*sua7-1*^ cells, thereby suggesting that initiation and elongation steps are not adversely affected in the mutant *in vivo* (Figure 2). Third, TFIIB physically interacts with the termination factors and this interaction is compromised in TFIIB^*sua7-1*^ mutant (Fig. 4). Fourth, TFIIB crosslinks to the 3’ end of genes in the TFIIB^*WT*^ cells but not in the termination defective TFIIB^*sua7-1*^ mutant (Singh and Hampsey, 2007).

TFIIB may affect termination either directly by interacting with termination factors and facilitating or stabilizing their recruitment at the 3’ end of genes or indirectly by influencing CTD-serine-2 phosphorylation. Our results favor a direct role for TFIIB in termination for two reasons. First, TFIIB crosslinks to the 3’ end of genes during transcription. Second, TFIIB exhibits physical interaction with subunits of CF1 and Rat1 termination complexes in the context of transcriptionally active chromatin. These interactions are compromised in the termination defective TFIIB^*sua7-1*^ mutant. The mammalian study demonstrating the TFIIB termination defect concluded that this defect observed in the absence of TFIIB was an indirect effect of an elongation defect due to excessive incorporation of P-TEFb on chromatin in the absence of TFIIB (Santana et al., 2022). However, the study never checked the presence of TFIIB at the 3’ end of genes or interaction of mammalian TFIIB with termination factors. An independent study, however, revealed that mammalian TFIIB, like its yeast homolog, also interacts with CstF-64, which is the mammalian homolog of yeast Rna15 (Wang et al., 2010). Furthermore, TFIIB-CstF-64 interaction was found to be crucial for termination of transcription in this study. These results corroborate the crucial role of TFIIB-termination factor interaction in termination of transcription. An independent study demonstrated the role of P-TEFb in regulating exoribonuclease activity of Xrn2, which is the mammalian counterpart of Rat1, in driving termination (Sanso et al., 2016). The interaction of TFIIB with Rat1 may similarly stimulate Rat1 activity leading to efficient termination.

Apart from the termination defect, the *sua7-1* mutant used in this study, also exhibits other phenotypes related to transcription. The transcription start site selection is altered in this mutant. The mutant is also defective in gene looping, which is the interaction of the promoter and terminator regions of a gene in a transcription-dependent manner (Singh and Hampsey, 2007). Previous results from our lab have shown that a gene loop is formed by the interaction of promoter-bound TFIIB with the termination factors occupying the 3’ end of genes (El Kaderi et al., 2009; Mayer et al., 2010; Medler et al., 2011). All gene looping defective mutants that we have analyzed so far in yeast are also defective in termination (Medler et al., 2011; Mukundan and Ansari, 2013; Al-Husini et al., 2013). Thus, TFIIB’s role in termination may be linked to its ability to facilitate gene loop formation. In fact, TFIIB-mediated gene looping has been found to affect alternative 3’ end processing, which is the selection of transcription termination site at the 3’ end of a gene in budding yeast (Lamas-Maceiras et al., 2016). A similar role of gene looping in alternative 3’ end processing was recently demonstrated in mammalian cells (Terrone et al., 2022). These results suggest that the role of TFIIB at the 3’ end of genes is not restricted to facilitating termination but may also involve selection of the transcription termination site from among multiple poly(A) sites present at the 3’ end (Al-Husini et al., 2020). Overall, this study serves as a paradigm for studying the broader role of TFIIB in the transcription cycle in higher eukaryotes. The study also opens up avenues for investigating the broader role of factors involved in different steps of the transcription cycle in the overall process of transcription and cotranscriptional RNA processing.

## Materials and Methods

### Yeast Strains

Yeast strains (*Saccharomyces cerevisiae*) used in this study are BY4733 with genetic background *MAT*α *his3*Δ*200* *trp1*Δ*63* *leu2*Δ*0* *met15*Δ*0* *ura3*Δ*0*. All subsequent strains used were derived from BY4733. Supplementary Material S1 lists all strains used in this study along with their genotype.

### Protein purification from soluble and chromatin fractions

Purification of TFIIB from soluble and chromatin fractions was performed exactly as described in O’Brien and Ansari (2024). Analysis was performed on four biological replicates, utilizing chromatin and soluble fractions obtained from *S. cerevisiae* strain *sua7-1-HA* (WA158; Supplemental Material S1).

### Mass spectrometry and quantitative analysis

Mass spectrometric analysis was performed as described in O’Brien and Ansari (2024). Analysis was performed with four biological replicates, utilizing chromatin and soluble fractions obtained from *S. cerevisiae* strain *sua7-1-HA* (WA158; Supplemental Material S1) in conjunction with *wild-type* data obtained in O’Brien and Ansari (2024).

### GRO-Seq

GRO-seq was performed essentially as described in O’Brien et al., (2023). Nascent, isolated RNA obtained using GRO-Seq was obtained from three biological replicates of *S. cerevisiae* strains *sua7-1* and *wild-type* (WA 304 and *BY4733*; Supplementary Material S1).

### GRO-Seq analysis

GRO-Seq samples were first stripped of adapter sequences using cutadapt. 3’ end reads were then aligned to the yeast s288c genome downloaded from SGD (version R64-3-1). To determine the readthrough index, we used the 3’ end annotations from SGD for mRNAs in YPD media. For each mRNA, we filtered the mRNAs to include only mRNAs whose 3’ end are at least 500 nucleotides from the next gene on the same strand. Next, we selected mRNAs that were at least 500 nucleotides in length or longer, and whose expression values were at least 1 read per nucleotide. We then calculated the readthrough index as the ratio of reads in a downstream window (50 to 500 nucleotides after the 3’ end) divided by an upstream window (-500 to -50 nucleotides in front of the 3’ end) (Baejen et al., 2017). To determine which mRNAs showed changes in readthrough index in the mutant (*sua7-1*), we compared the replicate readthrough index values from *WT* and mutant and used a 1-sided T-test (we ensured readthrough indices were measured in a minimum of five biological samples, with at least two from wild type and at least two in mutant). A criterion of a *p*-value less than 0.05 and a log2 fold change greater than 2 were used to determine which mRNAs showed termination defects.

To generate the metagene plot of the 3’ end, we filtered the mRNAs to include only mRNAs whose 3’ end are at least 500 nucleotides from the next gene on the same strand. Next, we selected mRNAs that were at least 500nt in length or longer, and whose expression values were at least 1 read/nt. For each mRNA, levels were normalized to the average read density in the CDS. The CDS normalized RNA levels were then averaged and plotted in a window from -200 nt before the 3’ end to +200 nt after the 3’ end. Outside lines represent standard error.

### TRO assay

Transcription Run-On assay was performed essentially as described in Dhoondia et al., (2017). Nascent, isolated RNA gathered using TRO was obtained from three biological replicates of *S. cerevisiae* strains *sua7-1* and *wild-type* (WA304 and *BY4733*; Supplementary Material S1). Primers used in TRO are described in Supplementary Material S2.

### ChIP

ChIP method and analysis was performed essentially as described in Dhoondia et al., (2021). Analysis was performed with three biological replicates, utilizing DNA obtained from ChIP of *S. cerevisiae* strains (Supplementary Material S1) harboring Pta1-HA (WA167/*wild-type* and WA383/*sua7-1*), Rat1-TAP (WA318/*wild-type* and WA384/*sua7-1*), and Rna15-TAP WA143/*wild-type* and WA386/*sua7-1*). Primers used in TRO are described in Supplementary Material S2.

## Data availability

The mass spectrometry proteomics data have been deposited to the ProteomeXchange Consortium via the PRIDE partner repository with the dataset identifier PXD041878 and 10.6019/PXD041878. Statistical source data is provided with this article. All other relevant data that support this study is available from the corresponding author upon reasonable request.

The GRO-Seq data has been deposited in NCBI database. The NCBI Geo accession number is: GSE259240.

## Acknowledgments

This work was supported by grants from National Institute of Health (1R01GM146803-05) and National Science Foundation (MCB1936030) to AA. We thank lab members Katie Dwyer, Emma Fidler, Travis Lenhausen, Hasan Jumani and Alden Kajy for useful help. We acknowledge the assistance of Dr. Paul Stemmer of Wayne State University Proteomics Core that is supported through National Institutes of Health grants P30 ES020957, P30 CA 022453 and S10 OD010700.

## References

1. Al-Husini, N., Kudla, P., & Ansari, A. (2013). A role for CF1A 3’ end processing complex in promoter-associated transcription. PLoS genetics, 9(8), e1003722.

2. Al-Husini, N., Medler, S., & Ansari, A. (2020). Crosstalk of promoter and terminator during RNA polymerase II transcription cycle. Biochimica et Biophysica Acta (BBA)-Gene Regulatory Mechanisms, 1863(12), 194657.

3. Allepuz-Fuster, P., O’Brien, M. J., González-Polo, N., Pereira, B., Dhoondia, Z., Ansari, A., & Calvo, O. (2019). RNA polymerase II plays an active role in the formation of gene loops through the Rpb4 subunit. Nucleic acids research, 47(17), 8975–8987.

4. Baejen, C., Andreani, J., Torkler, P., Battaglia, S., Schwalb, B., Lidschreiber, M., … & Cramer, P. (2017). Genome-wide analysis of RNA polymerase II termination at protein-coding genes. Molecular cell, 66(1), 38–49.

5. Chereji, R. V., Bharatula, V., Elfving, N., Blomberg, J., Larsson, M., Morozov, V., … & Björklund, S. (2017). Mediator binds to boundaries of chromosomal interaction domains and to proteins involved in DNA looping, RNA metabolism, chromatin remodeling, and actin assembly. Nucleic acids research, 45(15), 8806–8821.

6. Cho, E. J., & Buratowski, S. (1999). Evidence that transcription factor IIB is required for a post-assembly step in transcription initiation. Journal of Biological Chemistry, 274(36), 25807–25813.

7. Core, L. J., Waterfall, J. J., & Lis, J. T. (2008). Nascent RNA sequencing reveals widespread pausing and divergent initiation at human promoters. Science, 322(5909), 1845–1848.

8. Deng, W., & Roberts, S. G. (2005). A core promoter element downstream of the TATA box that is recognized by TFIIB. Genes & development, 19(20), 2418–2423.

9. Dhoondia, Z., Tarockoff, R., Alhusini, N., Medler, S., Agarwal, N., & Ansari, (2017). Analysis of termination of transcription using BrUTP-strand-specific transcription run-on (TRO) approach. JoVE (Journal of Visualized Experiments), (121), e55446.

10. El Kaderi, B., Medler, S., Raghunayakula, S., & Ansari, A. (2009). Gene looping is conferred by activator-dependent interaction of transcription initiation and termination machineries. Journal of Biological Chemistry, 284(37), 25015–25025.

11. Henriques T, Ji Z, Tan-Wong SM, Carmo AM, Tian B, et al. (2012) Transcription termination between polo and snap, two closely spaced tandem genes of D. melanogaster. Transcription 3: 198–212.

12. Lagrange, T., Kapanidis, A. N., Tang, H., Reinberg, D., & Ebright, R. H. (1998). New core promoter element in RNA polymerase II-dependent transcription: sequence-specific DNA binding by transcription factor IIB. Genes & development, 12(1), 34–44.

13. Lamas-Maceiras, M., Singh, B. N., Hampsey, M., & Freire-Picos, M. A. (2016). Promoter-terminator gene loops affect alternative 3’-end processing in yeast. Journal of Biological Chemistry, 291(17), 8960–8968.

14. Luse, D. S. (2014). The RNA polymerase II preinitiation complex: through what pathway is the complex assembled? Transcription, 5(1), e27050.

15. Mapendano, C.K., Lykke-Andersen, S., Kjems, J., Bertrand, E., Jensen, T.H. (2010) Crosstalk between mRNA 3’ end processing and transcription initiation, Mol Cell, 40, 410–422.

16. Mavrich, T. N., Ioshikhes, I. P., Venters, B. J., Jiang, C., Tomsho, L. P., Qi, J., … & Pugh, B. F. (2008). A barrier nucleosome model for statistical positioning of nucleosomes throughout the yeast genome. Genome research, 18(7), 1073–1083.

17. Mayer, A., Lidschreiber, M., Siebert, M., Leike, K., Söding, J., & Cramer, P. (2010). Uniform transitions of the general RNA polymerase II transcription complex. Nature structural & molecular biology, 17(10), 1272–1278.

18. Medler, S., Al Husini, N., Raghunayakula, S., Mukundan, B., Aldea, A., & Ansari, A. (2011). Evidence for a complex of transcription factor IIB (TFIIB) with Poly (A) polymerase and cleavage factor 1 subunits required for gene looping. Journal of Biological Chemistry, 286(39), 33709–33718.

19. Mischo, H. E., & Proudfoot, N. J. (2013). Disengaging polymerase: terminating RNA polymerase II transcription in budding yeast. Biochimica et Biophysica Acta (BBA)-Gene Regulatory Mechanisms, 1829(1), 174–185.

20. Mukundan, B., & Ansari, A. (2011). Novel Role for Mediator Complex Subunit Srb5/Med18 in Termination of Transcription. Journal of biological chemistry, 286(43), 37053–37057.

21. Murray, S. C., Serra Barros, A., Brown, D. A., Dudek, P., Ayling, J., & Mellor, J. (2012). A pre-initiation complex at the 3’-end of genes drives antisense transcription independent of divergent sense transcription. Nucleic acids research, 40(6), 2432–2444.

22. O’Brien, M. J., & Ansari, A. (2024). Protein interaction network revealed by quantitative proteomic analysis links TFIIB to multiple aspects of the transcription cycle. Biochimica et Biophysica Acta (BBA)-Proteins and Proteomics, 1872(1), 140968.

23. O’Brien, M. J., Gurdziel, K., & Ansari, A. (2023). Global Run-On sequencing to measure nascent transcription in Saccharomyces cerevisiae. Methods, 217, 18–26.

24. Paoletti, A. C., Parmely, T. J., Tomomori-Sato, C., Sato, S., Zhu, D., Conaway, R. C., … & Washburn, M. P. (2006). Quantitative proteomic analysis of distinct mammalian Mediator complexes using normalized spectral abundance factors. Proceedings of the National Academy of Sciences, 103(50), 18928–18933.

25. Pinto, I., Wu, W. H., Na, J. G., & Hampsey, M. (1994). Characterization of sua7 mutations defines a domain of TFIIB involved in transcription start site selection in yeast. Journal of Biological Chemistry, 269(48), 30569–30573.

26. Pugh, B. F., & Venters, B. J. (2016). Genomic organization of human transcription initiation complexes. PloS one, 11(2), e0149339.

27. Rhee, H. S., and B. F. Pugh. 2012. Genome-wide structure and organization of eukaryotic pre-initiation complexes. Nature 483:295–301.

28. Rossi, M. J., Kuntala, P. K., Lai, W. K., Yamada, N., Badjatia, N., Mittal, C., … & Pugh, B. F. (2021). A high-resolution protein architecture of the budding yeast genome. Nature, 592(7853), 309–314.

29. Roy, K., & Chanfreau, G. F. (2018). A global function for transcription factors in assisting RNA polymerase II termination. Transcription, 9(1), 41–46.

30. Sansó, M., Levin, R. S., Lipp, J. J., Wang, V. Y. F., Greifenberg, A. K., Quezada, E. M., … & Fisher, R. P. (2016). P-TEFb regulation of transcription termination factor Xrn2 revealed by a chemical genetic screen for Cdk9 substrates. Genes & development, 30(1), 117–131.

31. Santana, J. F., Collins, G. S., Parida, M., Luse, D. S., & Price, D. H. (2022). Differential dependencies of human RNA polymerase II promoters on TBP, TAF1, TFIIB and XPB. Nucleic acids research, 50(16), 9127–9148.

32. Singh, B. N., & Hampsey, M. (2007). A transcription-independent role for TFIIB in gene looping. Molecular cell, 27(5), 806–816.

33. Sun, Z. W., & Hampsey, M. (1996). Synthetic enhancement of a TFIIB defect by a mutation in SSU72, an essential yeast gene encoding a novel protein that affects transcription start site selection in vivo. Molecular and cellular biology, 16(4), 1557–1566.

34. Terrone, S., Valat, J., Fontrodona, N., Giraud, G., Claude, J. B., Combe, E., … & Bourgeois, C. F. (2022). RNA helicase-dependent gene looping impacts messenger RNA processing. Nucleic Acids Research, 50(16), 9226–9246.

35. Venters, B. J., & Pugh, B. F. (2013). Genomic organization of human transcription initiation complexes. Nature, 502(7469), 53–58.

36. Wang, Y., Fairley, J. A., & Roberts, S. G. (2010). Phosphorylation of TFIIB links transcription initiation and termination. Current Biology, 20(6), 548–553.

37. Woychik, N. A., & Hampsey, M. (2002). The RNA polymerase II machinery: structure illuminates function. Cell, 108(4), 453–463.

38. Wu, W. H., Pinto, I., Chen, B. S., & Hampsey, M. (1999). Mutational analysis of yeast TFIIB: a functional relationship between Ssu72 and Sub1/Tsp1 defined by allele-specific interactions with TFIIB. Genetics, 153(2), 643–652.

39. Zybailov, B., Mosley, A. L., Sardiu, M. E., Coleman, M. K., Florens, L., & Washburn, M. P. (2006). Statistical analysis of membrane proteome expression changes in Saccharomyces c erevisiae. Journal of proteome research, 5(9), 2339–2347.

