## Supplementary material for "Genome-wide analysis of TFIIB’s role in termination of transcription": Supplemnetary data

### S1 Yeast Strain Table

| Strain | Genotype |
|---|---|
| BY4733 | *MATα his3Δ200 trp1Δ63 leu2Δ0 met15Δ0 ura3Δ0* |
| WA143 | *BY4733, MATa his3Δ200 trp1Δ63 leu2Δ0 met15Δ0 ura3Δ0 RNA15-TAP, Histidine* marker |
| WA147 | *MATα his3Δ200 trp1Δ63 leu2Δ0 met15Δ0 ura3Δ0 SUA7-HA::HIS3 PCF11-Myc::TRP1* |
| WA158 | *MATα* cyc1-5000 cyc7-67 ura3-52 leu2-3,112 cyh2 *SUA7-1*-HA (KANMX) |
| WA167 | A HA-His cassette was inserted downstream ORF of *PTA1* in BEK1 strain (*MATα his3Δ200 trp1Δ63 leu2Δ0 met15Δ0 ura3Δ0* KANMX). |
| WA304 | *MATα his3Δ200 trp1Δ63 leu2Δ0 met15Δ0 ura3Δ0 sua7-1* |
| WA318 | *MATα his3Δ200 trp1Δ63 leu2Δ0 met15Δ0 ura3Δ0 RAT1-TAP Tryptophan* marker |
| WA383 | *MATα his3Δ200 trp1Δ63 leu2Δ0 met15Δ0 ura3Δ0 PARENT IS SUA7-1; PTA1-HA , Histidine* marker |
| WA384 | *MATα his3Δ200 trp1Δ63 leu2Δ0 met15Δ0 ura3Δ0 RAT1-TAP Tryptophan* marker, *Sua7-1* |
| WA386 | *MATα his3Δ200 trp1Δ63 leu2Δ0 met15Δ0 ura3Δ0 RNA15-TAP, Tryptophan* marker, *Sua7-1* |

### S2 Primer Table

***Primers used for TRO (RT-PCR):***

***18S:***

| Name | Sequence |
|---|---|
| 5' *18S* | GGAATAATAGAATAGGACGTTGG |
| 3' *18S* | GTTAAGGTCTCGTTCGTTATCG |

***BLM10:***

| Name | Sequence |
|---|---|
| *BLM10* 5'F | CCCTATGTTTTTCCCCTAC |
| *BLM10* 3'R | TCAGGCATAGTAACTTCTCCA |
| *BLM10* RT1 | ATCAGCAGATAGCTCCAG |
| *BLM10* RT2 | TGATTTATTACCCGGCAC |

***CBK1:***

| Name | Sequence |
|---|---|
| *CBK1* 5'F | ACATGGTGGTGCAGATG |
| *CBK1* 3'R | AATCTTCCTTGACTGGC |
| *CBK1* RT1 | AGAGGAATGTCCTTAACG |
| *CBK1* RT2 | AATGTGCGTTCTACATCC |
| *CBK1* RT3 | CCCTGATCTGCCATCTC |

***HEM3:***

| Name | Sequence |
|---|---|
| *HEM3* 5'F | TGTTCCGTTCCTATTGGTG |
| *HEM3* 3'R | AGAATTTTCTTTGCGCCAT |
| *HEM3* RT1 | ACAACGAAGCTTGTATGAATAG |
| *HEM3* RT2 | TTTTTAGAAGTTCACCAAGTG |

***KAP123:***

| Name | Sequence |
|---|---|
| *KAP123* 5'F | CTGTTCCAGCATTATTGG |
| *KAP123* 3'R | GAGCAACAATTCCGTTGT |
| *KAP123* RT1 | GTGCCTAAGATTGGGTGG |
| *KAP123* RT2 | CACATACATACAAGATTGGC |
| *KAP123* RT3 | TCATCGCTGAGTTGATCG |

| | |
|---|---|
| *KAP123* RT4 | CTTCGAGATGGACGATGC |
| *KAP123* RT5 | AATTGCGTCGTCAACATC |

***SEN1:***

| Name | Sequence |
|---|---|
| *SEN1* 5'F | GCAAATAGGACCCTCTCTC |
| *SEN1* 3'R | GGGATAAATGGGCTAGATGA |
| *SEN1* RT1 | GGCATTTCTGAAAGTGTAG |
| *SEN1* RT2 | ATTTAAAACAGCCCTCGAC |
| *SEN1* RT3 | CACTACCTTCTAAACCAATTG |

***SUR1:***

| Name | Sequence |
|---|---|
| *SUR1* 5'F | GCTTACTACAAGATGCATAG |
| *SUR1* 3'R | CCAGTGAGTACTTAGAAG |
| *SUR1* A1 | GTTGTGTATTGACGATACTG |
| *SUR1* A2 | ACGCAGGATAAGATGTGG |
| *SUR1* RT1 | GCAGTATCACTGGTACTT |
| *SUR1* RT2 | AGTCTGATATGAATTGTTGG |
| *SUR1* RT3 | ACAAGACTGGTCTTCAGG |
| *SUR1* RT4 | GTAGTATGTCGTAAGAAGC |
| *SUR1* RT5 | GTATAGAGGGTAATATCACAG |

***Primers used for ChIP:***

***BLM10:***

***Pta1: 1:****S/T3R,* ***2:****T1/T3R,* ***3:****T1/T6R,* ***4:****T2/T6,* ***5:****T6/T8*

***Rna15: 1:****S/T3R,* ***2:****T1/T3R,* ***3:****T4/T6R*

***Rat1: 1:****S/T3R,* ***2:****T1/T6R,* ***3:****T2/T6R,* ***4:****T2/T8R,* ***5:****T4/T6R*

***Ser2p: 1:****S/T2R,* ***2:****T2/T6R,* ***3:****T2/T8R,* ***4:****T4/T6R,* ***5:****T6/T8R*

***Ser5p 5' End: 1:****AF/CR,* ***2:****BF/BR,* ***3:****BF/CR,* ***4:****CF/CR*

***Ser5p 3' End: 1:****S/T3R,* ***2:****T1/T3R,* ***3:****T1/ T6R,* ***4:****T2/T6R,* ***5:****T2/ T8R,* ***6:****T4/T6R*

| Name | Sequence |
|---|---|
| *BLM10* S | ATATCAGCGTTCCCCTATG |
| *BLM10* T1 | ATGGAGAAGTTACTATGCC |
| *BLM10* T2 | ACTTGGAAGTTCGATAGAG |
| *BLM10* T4 | TAGTGTAACTATGATCACGGC |
| *BLM10* T6 | GCATTCTACCTCTAGGGAAG |

| | |
|---|---|
| *BLM10* T2R | CTGGAGCTATCTGCTGAT |
| *BLM10* T3R | GTGCCGGGTAATAAATCAA |
| *BLM10* T6R | CTGTATATACACGAGGGCG |
| *BLM10* T8R | GGAAAGAGCTATGACTCTAT |
| *BLM10* AF | GACGATGATATCAAATCACCC |
| *BLM10* BF | GAGAGAAGTCCAGGAAGG |
| *BLM10* CF | ACGCTGGATTATGTTAGTGAC |
| *BLM10* AR | CCTATTGAAGAGTTCTTACCA |
| *BLM10* CR | GACTTACTTGGTAACGATATTGCT |

***HEM3:***

***Pta1: 1:****MF/MR,* ***2:****5F/3R,* ***3:****5F/4R,* ***4:****7F/7R,* ***5:****7F/8R*

***Rna15: 1:****MF/MR,* ***2:****5F/4R,* ***3:****5F/4AR,* ***4:****6F/8R,* ***5:****7F/6R*

***Rat1: 1:****MF/MR,* ***2:****5F/4R,* ***3:****5F/4AR,* ***4:****6F/4R,* ***5:****6F/5R*

***Ser2p: 1:****M/FMR,* ***2:****7F/8R,* ***3:****8F/6R,* ***4:****8F/8R,* ***5:****8F/9R*

***Ser5p 5' End: 1:****A/AR,* ***2:****A/BR,* ***3:****A/CR,* ***4:****B/AR*

***Ser5p 3' End: 1:****MF/MR,* ***2:****6F/8R,* ***3:****7F/8R,* ***4:****7F/9R,* ***5:****8F/6R*

| Name | Sequence |
|---|---|
| *HEM3* MF | AGAAAGGGTGACACCAAGATG |
| *HEM3* 5F | CACCATTAATTTGATCAGTCC |
| *HEM3* 6F | CCTTATCTCCTTTATCTTTCAGT |
| *HEM3* 7F | GACCATTTTCAAATGACCAC |
| *HEM3* 8F | ACTATTCATACAAGCTTCGTTG |
| *HEM3* MR | GTTGACGTTGAAGGCACAGAA |
| *HEM3* 3R | CCCTCTCATTCCTCTCAT |
| *HEM* 4R | CAAATGACCACTGCATAAATAT |
| *HEM* 4AR | GAATATGTAATTCCTACCTTTG |
| *HEM3* 5R | CTATTCATACAAGCTTCGTTGT |
| *HEM3* 6R | CACTTGGTGAACTTCTAAAAA |
| *HEM3* 8R | GATCAGAATTCCGACTCCAG |
| *HEM3* 9R | CTATCTCTGCCACTCACTAT |
| *HEM3* AF | GGTGGGAGAAAATCGAAATTG |
| *HEM3* BF | CTGATCGAAGAAAAGTATCCG |
| *HEM3* AR | CCAACAGATTGTCTTGTCATG |
| *HEM3* BR | GTCTCTGGATGACCTTCCA |
| *HEM3* CR | GACACCAAGATGATGAAGATTC |

***KAP123:***

***Pta1: 1:****F2/R1,* ***2:****DN1/R7,* ***3:****DN2/R8,* ***4:****DN3/DN10,* ***5:****DN4/DN10,* ***6:****DN5/DN13*

***Rna15: 1:****F2/R1,* ***2:****DN1/R7,* ***3:****DN2/R8,* ***4:****DN3/DN10,* ***5:****DN4/DN10,* ***6:****DN5/DN13*

***Rat1:*** *1:F2/R1,* ***2:****DN3/DN10,* ***3:****DN4/DN10,* ***4:****DN5/DN12,* ***5:****DN5/DN13,* ***6:****DN6/DN12*

***Ser2p: 1:****DN1/R7,* ***2:****DN4/DN11****,*** ***3:****DN5/DN11,* ***4:****DN5/DN12,* ***5:****DN6/DN12*

***Ser5p 5' End: 1:****F1/R3,* ***2:****F2/R3,* ***3:****F3/R1,* ***4:****F4/R1*

***Ser5p 3' End: 1:****DN3/DN10,* ***2:****DN4/DN11,* ***3:****DN5/DN11*

| Name | Sequence |
|---|---|
| *KAP123* F1 | CTGTTCCAGCATTATTGG |
| *KAP123* F2 | GTGCTCTAGTTCCATTGG |
| *KAP123* F3 | TGCTAGTGGATGTGTGG |
| *KAP123* F4 | TTAGCAGCCGAAGATGAC |
| *KAP123* R1 | CCTTATGCTTCATCTCTTC |
| *KAP123* R3 | GTGCCTAAGATTGGGTGG |
| *KAP123* R7 | AATTGCGTCGTCAACATC |
| *KAP123* R8 | GTTCTGGGATATTCGCCG |
| *KAP123* DN1 | CACCCAATCTTAGGCAC |
| *KAP123* DN2 | GCCAATCTTGTATGTATGTG |
| *KAP123* DN3 | GTTGACGACGCAATTCG |
| *KAP123* DN4 | GCAAGTTCAGGTCATCTTG |
| *KAP123* DN5 | TTGACGATGTATAGTACATG |
| *KAP123* DN6 | TGCAAGTTACCTGCAGG |
| *KAP123* DN10 | CCTGCAGGTAACTTGC |
| *KAP123* DN12 | TTGAACTGTGGTCCCTG |
| *KAP123* DN13 | GCACTATACACTCGTTGG |

***SUR1:***

***Pta1: 1:****A1/R1,* ***2:****A1/R2,* ***3:****F0.5/R1,* ***4:****DN1/R5,* ***5:****DN3/R1*

***Rna15: 1:****F2/A2,* ***2:****F0.5/R1,* ***3:****DN1/R5,* ***4:****DN2/R6,* ***5:****DN3/R7,* ***6:****DN6/DN11*

***Rat1: 1:****F2/A2,* ***2:****F0.5/R1,* ***3:****DN1/R5,* ***4:****DN2/R6,* ***5:****DN3/R7,* ***6:****DN6/DN11*

***Ser2p: 1:****F3/A2, 2:DN1/R5,* ***3:****DN2/R6,* ***4:****DN3/R7,* ***5:****DN5/DN11*

***Ser5p 5' End: 1:****A2F/AF1,* ***2:****F7/A2R,* ***3:****F3/A2R,* ***4:****F3/A3R,* ***5:****F1/A2R*

***Ser5p 3' End: 1:****F3/A2,* ***2:****DN1/R5,* ***3:****DN7/DN12*

| Name | Sequence |
|---|---|
| *SUR1* A1 | GCTTACTACAAGATGCATAG |

| | |
|---|---|
| *SUR1* A2 | ATGTCTCGATCTACATCC |
| *SUR1* F0.5 | CCACATCTTATCCTGCG |
| *SUR1* F1 | GCTTACTACAAGATGCATAG |
| *SUR1* F2 | GCGCATACCTAAGAACG |
| *SUR1* F3 | CCCTTACATGACTATTATGG |
| *SUR1* F7 | CATAGAACGTGCCGATG |
| *SUR1* R1 | GCAGTATCACTGGTACTT |
| *SUR1* R2 | CCAGTGAGTACTTAGAAG |
| *SUR1* R5 | GTAGTATGTCGTAAGAAGC |
| *SUR1* R6 | ATGATTATGTTGATGCTGG |
| *SUR1* R7 | GTATAGAGGGTAATATCACAG |
| *SUR1* DN1 | CTCCAACAATTCATATCAGAC |
| *SUR1* DN2 | CTGAAGACCAGTCTTGTAC |
| *SUR1* DN3 | CTTCTTACGACATACTACTCC |
| *SUR1 DN5* | ATTACCCTCTATACTACGATATC |
| *SUR1* DN6 | GAATATGAACACTTCCTCAC |
| *SUR1* DN7 | CATCGCAACGGAGTAAG |
| *SUR1* DN11 | GCGGCTACTAGTCGAG |
| *SUR1* A1F | GTTGTGTATTGACGATACTG |
| *SUR1* A1R | ATCGATGTATACACCACC |
| *SUR1* A2R | ACGCAGGATAAGATGTGG |
| *SUR1* A3R | ACGCAGGATAAGATGTGG |

### S3 Supplementary Figure 1

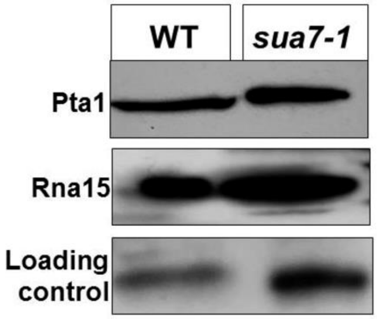


***Supplementary Figure 1: Termination factor abundance is not reduced in sua7-1 mutant.*** *Western blot depicting the detection of termination factors Pta1 and Rna15, compared between wild-type and the sua7-1 mutant, along with loading control.*

### S4 Supplementary Figure 2

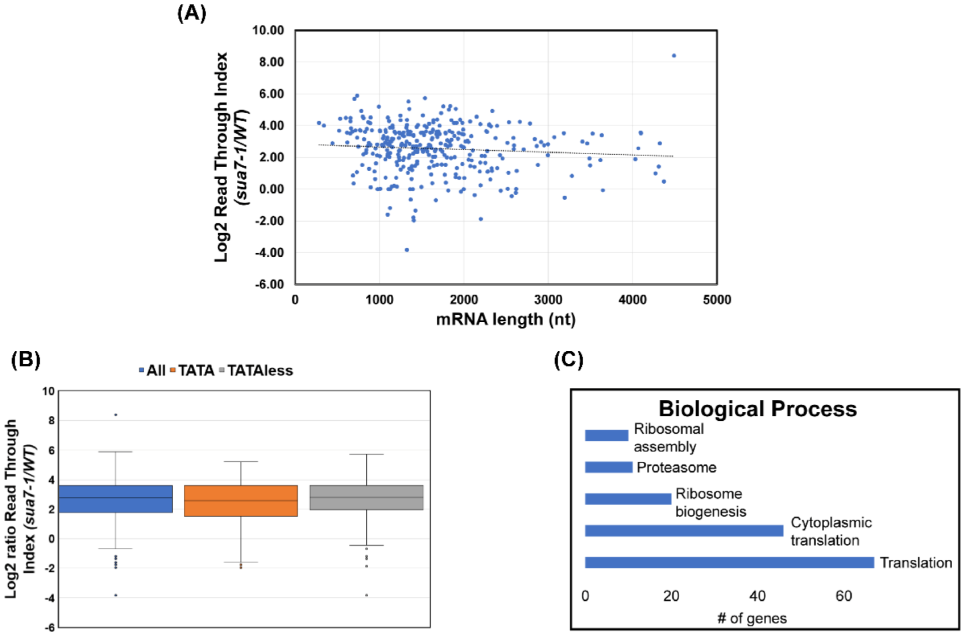


***Supplementary Figure 2: TFIIB-dependent termination defect is not correlated to gene length, is independent of TATA box presence in promoter, and is pronounced in translation as well as ribosome function related genes. (A)** mRNA length is plotted on the x-axis and log2 RTI ratio of sua7-1/WT is on y=axis. A trendline is shown as a dotted line. Data is from three biological replicates. **(B)** The median log2 ratio of RTI for sua7-1/WT is compared across total genes analyzed (blue), genes with an associated TATA box (orange), and TATA-less genes (grey). Within the box and whisker plot, the middle line represents the median value of the corresponding RTI. The top and bottom lines represent the upper and lower quartiles. Data is from three biological replicates. Gene ontology analysis for total analyzed genes. The number of genes (x-axis) and their corresponding biological process (y-axis) are shown. Terms and analysis performed with NCBI DAVID gene ontology database.*

### S5 Supplementary Figure 3

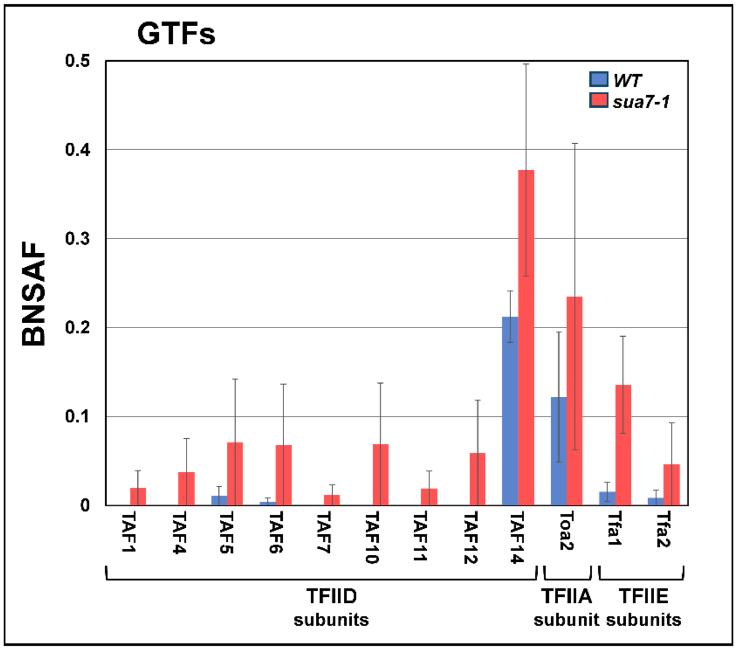


***Supplementary Figure 3: TFIIB-general transcription factors interactions are not compromised in sua7-1 mutant.*** *BNSAF values of subunits of TFIID, TFIIA and TFIIE in affinity purified TFIIB from chromatin fraction in sua7-1 and WT cells. Data is form at least three biological replicates. p-values obtained from a standard, paired t-test. Error bars represent one unit of standard error based on four biological replicates.*
